# Gene conversion limits the cost of asexuality in somatically immortal worms

**DOI:** 10.1101/2023.03.20.533553

**Authors:** Simon Kershenbaum, Manuel Jara-Espejo, Ashleigh S. Griffin, Aziz Aboobaker

## Abstract

Most multicellular organisms reproduce sexually despite the costs associated with sexuality. This has been explained as the result of selection favouring the ability to recombine the genome. The lack of recombination in asexual species constrains their adaptability and leads to the accumulation of deleterious mutations, ultimately increasing their risk of extinction. Nonetheless, successful asexual life histories persist among multicellular organisms, and explanatory mechanisms which may help limit the cost of asexuality remain enigmatic. In search of these mechanisms, we looked at that the molecular evolutionary changes in the sexual and obligately asexual strains of the planarian flatworm, *Schmidtea mediterranea*. We find that the accumulation of deleterious mutations in highly conserved genes is largely avoided in the asexual strain. We find evidence that this is achieved by somatic gene conversion in stem cells allowing for the restoration of beneficial alleles and purification of deleterious mutations. Taken together, our analysis identifies gene conversion as a mechanism which may contribute to the maintenance of asexuality in an obligately fissiparous metazoan.

## Introduction

Most multicellular life reproduces sexually, while obligate asexuality is exceedingly rare (Judson and Normark, 1996). This pattern persists despite the high costs of sex, including the “two-fold” cost of sex (Gibson et al., 2017), the breaking up of beneficial haplotypes (Charlesworth and Barton, 1996), and the broad costs associated with securing a mate (Lehtonen et al., 2012). The maintenance of sexuality in the face of such costs suggests that asexuality, the alternative reproductive strategy, is somehow fundamentally limited and is outcompeted by sexual strategies in the long term (Lynch et al., 1993). However, some ancient asexual organisms exist, which raises the question of how they avoid the limitations of asexuality (Bast et al., 2018; Butlin et al., 1998; Judson and Normark, 1996; Lázaro et al., 2011; Wilson et al., 2018).

The potential disadvantages of asexual reproduction are driven by the tight linkage between alleles in an asexual genome (Box. 1) (Comeron et al., 2008; Felsenstein, 1974; Hill and Robertson, 1966). This linkage means that selection cannot act on individual alleles or allele combinations, but instead must act on the genomes of individuals as a whole - a problem known as “The Hill-Robertson effect” (Comeron et al., 2008; de Visser and Elena, 2007; Hill and Robertson, 1966). This reduces the adaptability of asexual populations, which has been shown to be particularly problematic in the context of changing environments and biotic competition (Hamilton et al., 1990; Lively and Morran, 2014; Vergara et al., 2014).

Additionally, the inability to remove deleterious mutations from the next generation through recombination results in the accumulation of deleterious mutations over time in a process called “Muller’s ratchet” (Lynch et al., 1993; Muller). This occurs since even the fittest individual in an asexual population contains some deleterious mutations, and as this fittest genome reaches fixation, so do its deleterious mutations, leading to a ratchet-like process of mutation accumulation (Lynch et al., 1993; Muller). Taken together, the reduced ability to unlink alleles in asexual genomes is thought to drive Muller’s ratchet and the Hill-Robertson effect, both suggested as explanations for why obligate asexual reproduction is rare (Felsenstein, 1974).

### Box. 1

The limitations of obligate asexual reproduction stem from the tight linkage of alleles in the genomes that lack recombination. Muller’s ratchet describes how this lack of recombination leads to the accumulation of deleterious mutations over time. This is because even the fittest asexual genome will have some deleterious mutations that will be passed on to the next generation, while in sexual reproduction, genomes that only contain beneficial alleles can theoretically be generated and selected for. Hill-Robertson interference is a broad term describing the inefficient selection that occurs when alleles are tightly linked due to the lack of recombination. This means that deleterious mutations can interfere with the selection for beneficial mutations (and vice versa), while in sexual populations these mutations can be selected upon independently. Clonal interference describes the inefficacy resulting from the fact that two beneficial mutations in different lineages will compete with one another. Since asexual lineages don’t recombine with one another, only the fittest variant will persist, while in sexual populations a genome containing both variants can be formed.

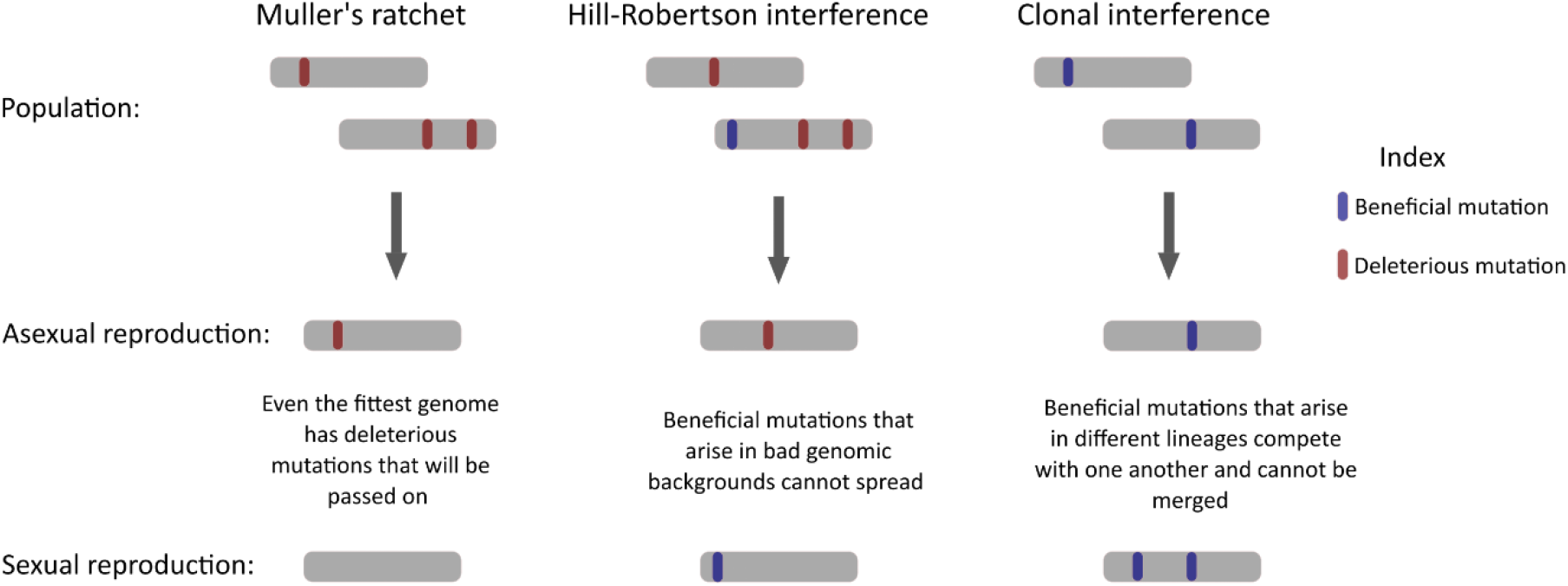

However, some obligate asexual species persist over evolutionary time (Barraclough et al., 2007; Bast et al., 2018; Lázaro et al., 2011; Schön et al., 2009), suggesting that they may have evolved mechanisms to limit the impact of long-term asexuality. Such ancient asexuals include bdelloid rotifers, oribatid mites, darwinulid ostracods and more, all of which have likely been asexual for millions of generations (Brandt et al., 2017; Brandt et al., 2021; Flot et al., 2013; Mark Welch and Meselson, 2000; Schön et al., 2003; Schön et al., 2009). To date, there is no consensus regarding the methods by which they avoid the costs associated with asexuality. One suggested mechanism is gene conversion (Box. 2): a form of mitotic recombination in which one allele overwrites another (Butlin et al., 1998; Flot et al., 2013; Schön et al., 2009). While deleterious and beneficial alleles are equally likely to get overwritten in the process of gene conversion, selection may be able to act to remove the deleterious gene conversion events and drive the spread of beneficial alleles (Box. 2).

Ancient asexuals such as bdelloid rotifers (Flot et al., 2013) and darwinulid ostracods (Schön et al., 2009) exhibit excess homozygosity, which has been proposed to be the result of gene conversion, leading the authors to suggest a role for gene conversion in the long term survival of these asexuals. However, evidence from other species suggests that gene conversion may not be sufficient to limit mutation accumulation, since young asexual lineages of Daphnia show relatively high rates of gene conversion while still showing a significant accumulation of deleterious mutations (Omilian et al., 2006; Tucker et al., 2013). In neither case has it been shown that selection acts upon gene conversion events to remove deleterious mutations and fix beneficial alleles in the population. Taken together, there is currently no consensus about the mechanisms that limit the costs of asexuality in ancient asexuals, and we need to assess more groups of long-term asexuals with diverse modes of asexual reproduction to better understand how ancient asexuals might overcome the limitations of asexuality.

The planarian flatworm *S. mediterranea* is a highly regenerative animal that has both a sexual and an asexual strain (Tan et al., 2012). The sexual strain is a cross fertilizing hermaphrodite and the obligate asexual strain lacks reproductive organs, reproducing solely by fission followed by whole body regeneration (Newmark et al., 2008; Saberi et al., 2016) (Fig. 1).

**Figure 1.**
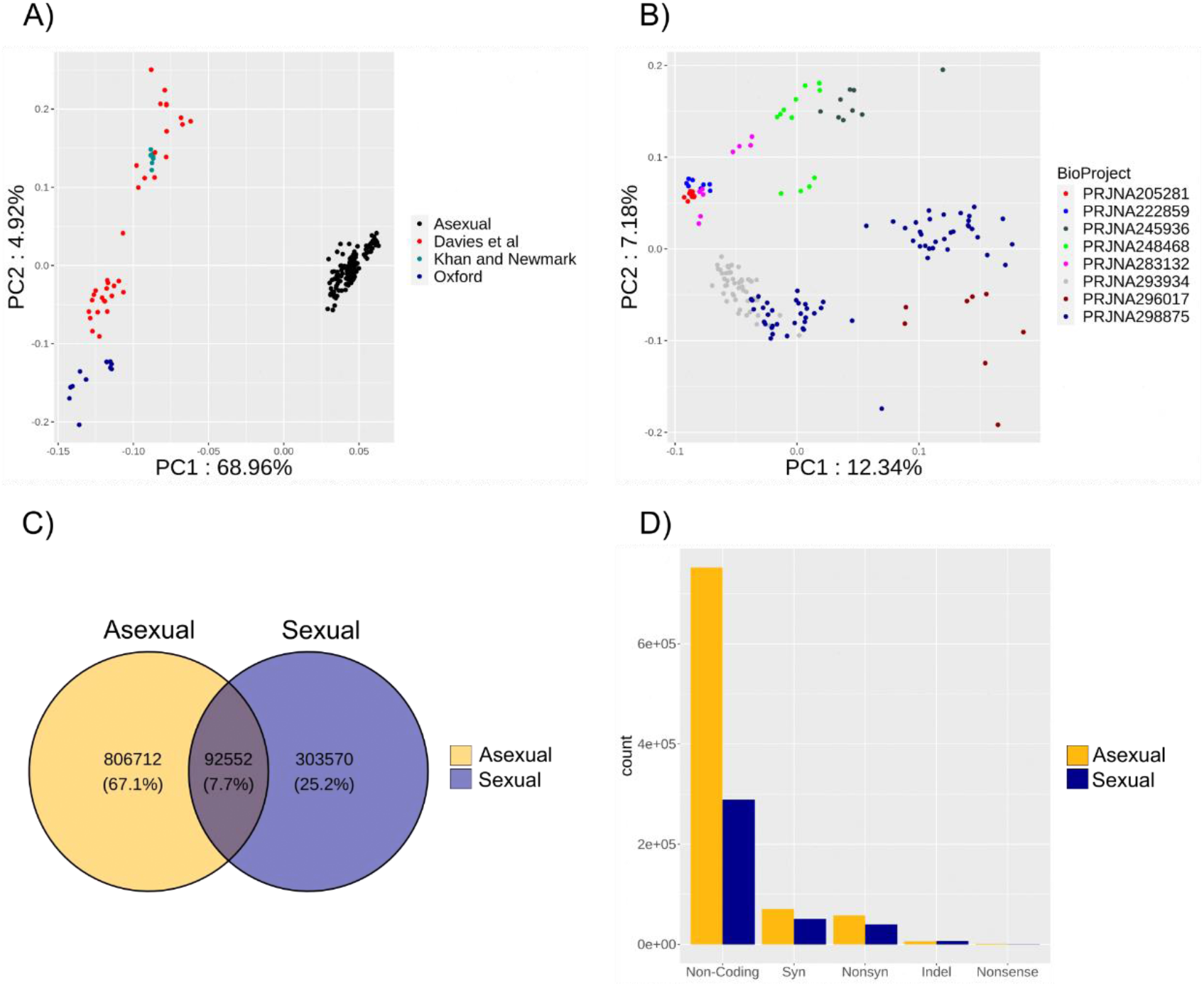
- Description of population structure and variants. A) PCA of all libraries in study; B) PCA of asexual libraries in study coloured by BioProject ID; C) Venn Diagram showing the percentage of shared and unique variants between sexual and asexual populations. D) Barplot showing the numbers of different types of variants in sexual and asexual populations.

The asexual strain is evidently able to thrive, despite the limitations of asexual reproduction, providing us with an opportunity to examine potential mechanisms for dealing with the limitations of asexuality (Lázaro et al., 2011; Sahu et al., 2017; Tan et al., 2012). The fissiparous nature of asexual planarian reproduction is based on whole body regeneration that is mediated by somatic, immortal, pluripotent adult stem cells called neoblasts (Wagner et al., 2011). Continuous BrdU labelling suggests that these neoblasts have a generation time of 24- 72 hours as they are constantly in the cell cycle, suggesting that since the origin of the asexual lineage there have been at least tens to hundreds of millions of asexual neoblast generations (Fields and Levin, 2018; Newmark and Sánchez Alvarado, 2000), dependent upon when the asexual strain arose (Lázaro et al., 2011). Such a high number of asexual generations is equivalent to some of the previously studied ancient asexuals but represents an understudied mode of asexual reproduction in metazoans. This fissiparous lifestyle and the presence of a closely related sexual strain which allows for the direct genetic comparison of sexual and asexual strains, makes *S. mediterranea* a valuable species for understanding how such asexual lineages might persist over evolutionary time.

In this study, we investigate how asexual planarians may limit the costs of asexuality. Specifically, we test whether planarian flatworms are able to escape the costs of asexuality through gene conversion. We find that asexual planarians limit the impact of Muller’s ratchet, and that they show GC-biased excess homozygosity indicative of gene conversion.

Furthermore, we find that mutations that are predicted to be more deleterious are less homozygous, suggesting that gene conversion events can be acted upon by selection to fix the beneficial allele in the population and remove deleterious mutations. Thus, we find evidence that gene conversion mediated removal of deleterious mutations may allow asexual planarians to limit Muller’s ratchet and persist despite their obligate asexuality.

## Results

### Population structure of laboratory *S. mediterranea*

To better understand how laboratory populations of S. mediterranea are related to one another, we used 169 asexual RNAseq libraries from 7 studies and 46 RNAseq libraries from 3 studies and evaluated their population structure (SI1) (Currie and Pearson, 2013; Davies et al., 2017; Duncan et al., 2015; Khan and Newmark, 2022; Scimone et al., 2014; Tu et al., 2015; van Wolfswinkel et al., 2014; Vogg et al., 2014; Zhu et al., 2015). We found that all asexual samples clustered together independently from the sexual samples (Fig. 1A). Asexual populations also largely clustered by BioProject, suggesting that populations are evolving independently to a certain extent in each laboratory (Fig. 1B). To ensure that no one population would be biasing the results, for example by being abnormally homozygous, we checked for outliers in mean allele frequency and proportion of homozygosity across all BioProjects (Fig. S1). We found no outliers in the asexual population (Fig. S1A-C). One sexual population(Davies et al., 2017), had a higher proportion of rare alleles compared to the other sexual populations (Fig. S1D). To ensure that our results are not biased by this population, all further analyses comparing asexual to sexual allele frequencies were also done separately against each sexual population. Most of the variants isolated were specific to the asexual lineage (806,712 variants, which amounts to 67.1% of the variants found in this study), and only 7.7% were shared between the sexual and asexual populations (Fig. 1C, Fig. S1E). This abundance of asexual variants is expected since variants were called against the available sexual genome (Grohme et al., 2018). In both sexual and asexual populations, the majority of variants were non-coding, followed by synonymous mutations, non-synonymous mutations, and indels respectively (Fig. 1D, Fig. S1F). In summary, our analysis suggests that asexual populations form subpopulations across different laboratories, and are distinct from the sexual population, with over 800,000 variants that are unique to the asexual population.

### Limited impact of Muller’s ratchet in asexual planarians

We began by looking for evidence of deleterious mutation accumulation in the obligate asexual strain by using dN/dS analyses (SI2-3). We found that across the entire set of protein coding genes (Neiro et al., 2022) (SI4), the median asexual dN/dS value is 3.6% higher than that found in the sexual strain (Wilcoxon rank sum; NAsexual = 16,445, NSexual = 10,700, W = 91,647,379, P < 0.001, Fig.2A). High dN/dS values can indicate both deleterious mutation accumulation and positive selection, thus we assessed dN/dS values in genes that are unlikely to be under positive selection. Specifically, we compared the dN/dS values of exons belonging to highly conserved single copy BUSCO genes (SI5) (Manni et al., 2021), and found no significant difference between the strains (Wilcoxon signed rank; Npairs = 160, V = 7,069, P = 0.284, Fig. 2B). However, when we compared the dN/dS values of exons under significant negative selection in both strains (SI6), we found that the asexual dN/dS values were 27% higher than in the sexual strain (Wilcoxon signed rank; Npairs = 41, V = 665, p-value = 0.002, Fig. 2C). This is in contrast to some previous studies that found higher impacts of Muller’s ratchet in asexual lineages (dN/dS values ∼200%-∼3,000% higher than in their sexual counterparts) (Barraclough et al., 2007; Bast et al., 2018; Henry et al., 2012; Johnson and Howard, 2007; Neiman et al., 2010; Paland and Lynch, 2006). Additionally, we isolated genes that contain exons under positive selection in the asexual population (Fig. 2D, SI7-8). These included multiple genes related to GLIPR1-like genes (see methods), which may suggest that these genes evolved new functions in the asexual population. Taken together, while asexual planarians may be subject to Muller’s ratchet, the accumulation of deleterious mutations is somehow limited, and is not present in highly conserved single copy genes. This requires a mechanistic explanation in the absence of sexual recombination.

**Figure 2.**
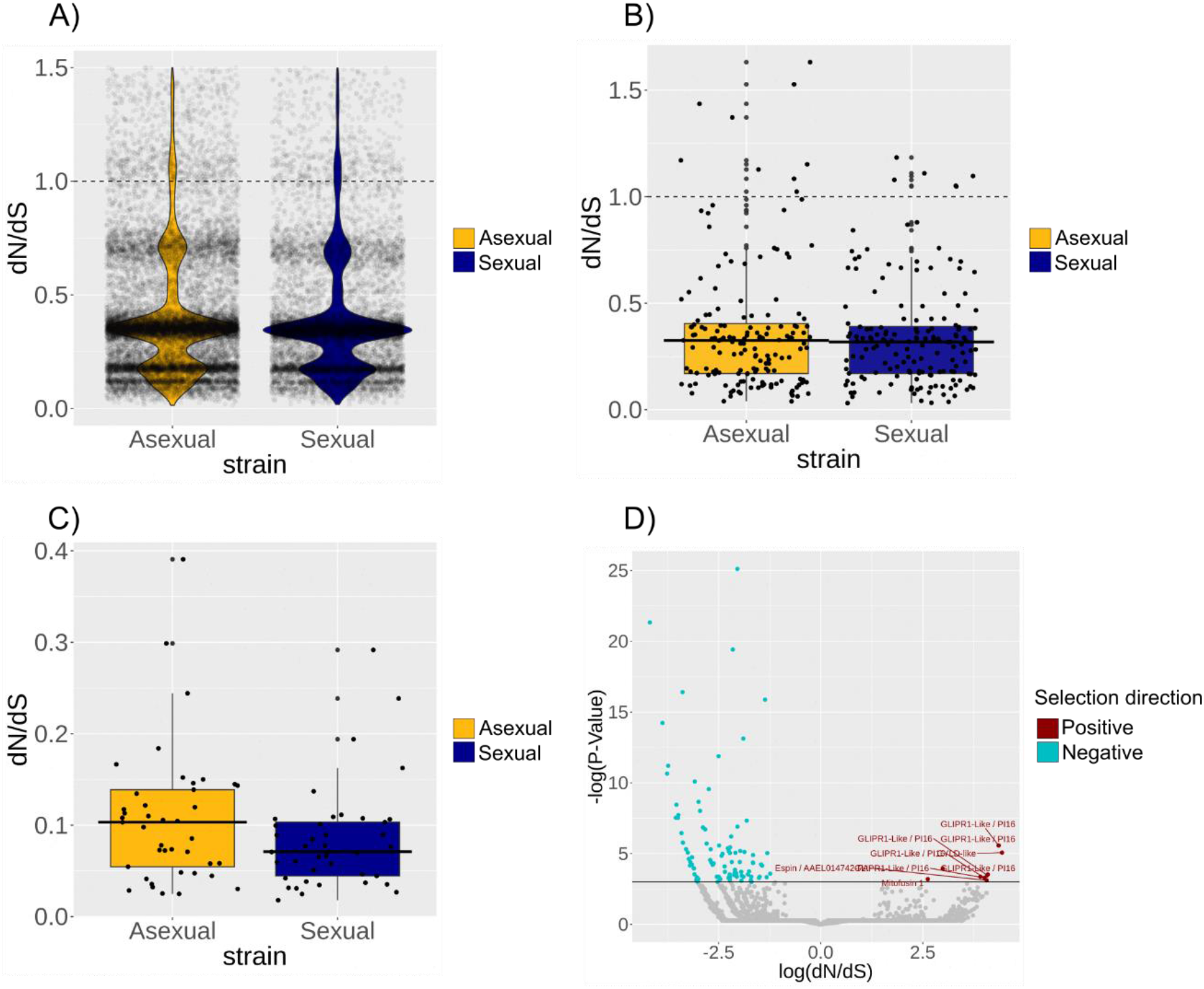
- dN/dS analysis in sexual and asexual populations. A) Violin plot showing the dN/dS values of all exons for both sexual and asexual populations. Sexual mean dN/dS = 0.444, asexual mean dN/dS = 0.46 ; B) dN/dS of all BUSCO exons that contained both synonymous and nonsynonymous variants in both sexual and asexual population. Sexual mean dN/dS = 0.318, asexual mean dN/dS = 0.325; C) dN/dS of exons under negative selection in both sexual and asexual populations. Sexual mean dN/dS = 0.084, asexual mean dN/dS = 0.107; D) Volcano plot showing exons under positive selection in the asexual population.

### Loss of heterozygosity mechanisms in asexuals may limit Muller’s ratchet

Mechanisms limiting Muller’s ratchet in asexual planarians may involve somatic recombination mechanisms, such as gene conversion, that in combination with selection can remove deleterious mutations (Box. 2). A signature of such a process would be homogenisation of the genome, leading to low polymorphism and high homozygosity. We compared allele frequencies of synonymous variants in asexual and sexual populations and found that asexual allele frequency is significantly higher than in the sexual population for both shared ancestral variants and lineage specific variants suggesting that there is lower polymorphism in the asexual population (Wilcoxon signed rankancestral; Npairs = 9,206, V = 40,369,398, Pancestral < 0.001, Fig. 3A; Wilcoxon rank sumlineage-specific, Nasexual = 30,463, Nsexual = 13,428, W = 338,332,851, Plineage-specific < 0.001, Fig. 3B). Additionally, in asexuals, the allele frequencies of ancestral variants were 45% higher than lineage specific variants, while in the sexual population the allele frequencies of ancestral variants were only 4% higher than lineage specific variants (Wilcoxon rank sum; Nasexual = 309, Nsexual = 305, W = 82,102, P < 0.001, Fig. 3C). The increased allele frequency in ancestral variants suggests a process of homogenisation over time, which is more effective in the asexual genome.

**Figure 3.**
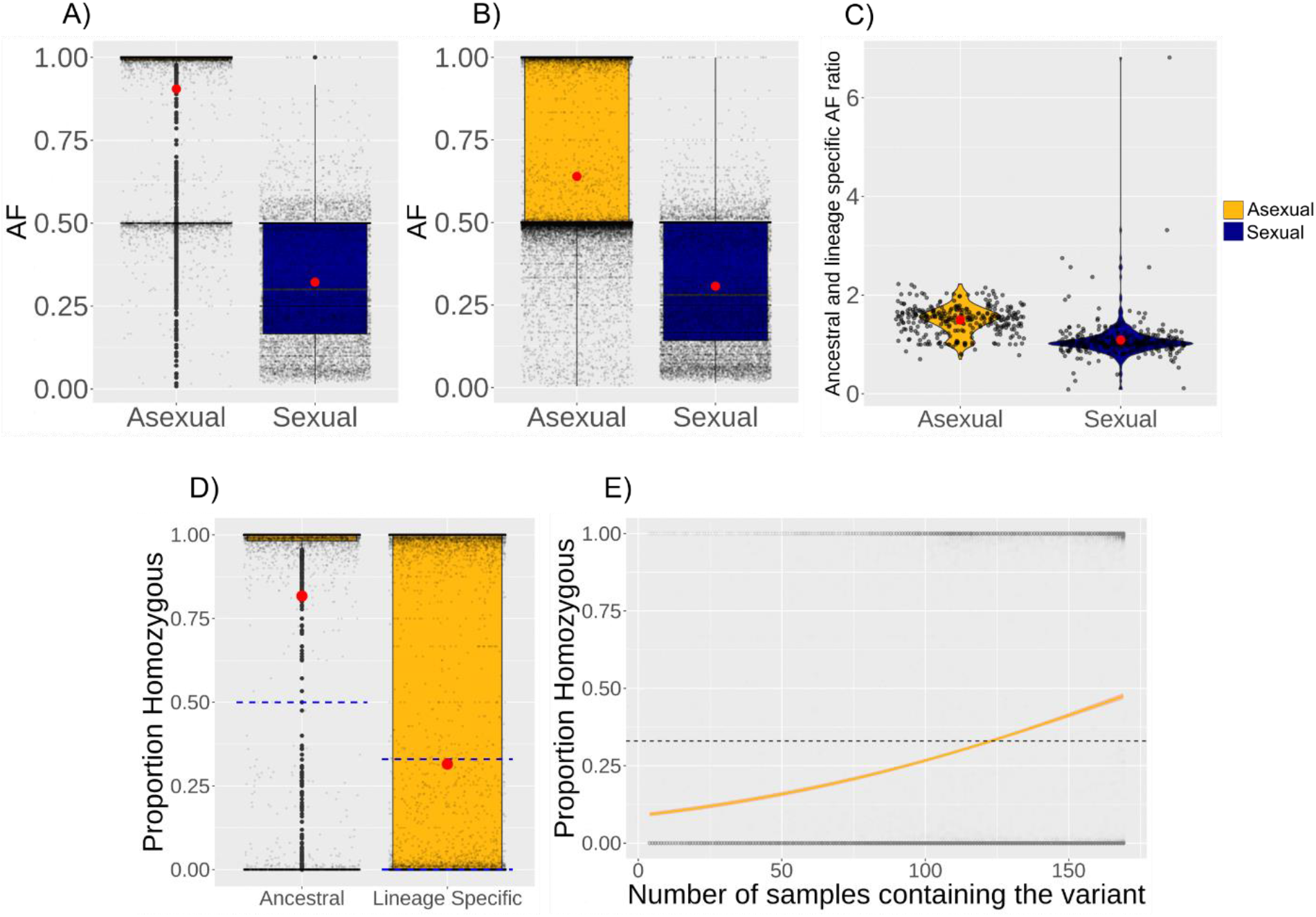
- Excess homozygosity in the asexual population. Red dots indicate means. A) Allele frequencies of ancestral variants shared by sexual and asexual populations. Mean sexual allele frequency = 0.32. Mean asexual allele frequency = 0.9; B) Allele frequencies of lineage specific. Mean sexual allele frequency = 0.31. Mean asexual allele frequency = 0.64; C) Violin plot showing the ratio of average allele frequencies per scaffold of ancestral vs lineage specific variants. Mean sexual ratio = 1.038. Mean asexual ratio= 1.446; D) Boxplot showing the proportion of homozygous libraries per variant in ancestral (shared between the sexual and asexual populations) and lineage specific variants in the asexual population. Dashed blue lines indicate the expected proportion of homozygosity in the absence of recombination; E) Binomial GLM showing that the proportion of homozygosity in asexual lineage specific variants increases with the number of libraries containing the variant. When all libraries contain the variant the mean proportion of homozygosity is 0.58. The dashed line indicates the expected homozygosity assuming that all asexual specific variants were inherited from sexual ancestors.

High allele frequencies in asexuals may be explained by the clonal nature of asexual planarians, so we tested whether they also show excess homozygosity which would be indicative of a homogenising process that cannot be accounted for by clonality. In the absence of recombination, half of the ancestral variants shared between the sexual and asexual strains should be homozygous in the asexual strain (see methods). However, we find that the mean proportion of homozygosity in asexual ancestral variants is 0.82, suggesting that some homogenising process is taking place over time (Wilcoxon signed rank; Nasexual- ancestral = 9,206, V = 34,205,206, P < 0.001, Fig. 3D).

We also tested whether there is excess homozygosity in the uniquely asexual variants. For asexual specific variants, the expected proportion of homozygosity depends on the proportion of variants that formed by mutation after the sexual-asexual split, versus those that were inherited from their sexual ancestors and subsequently lost from the sexual population. At the extremes, if all of them formed after the sexual-asexual split we would expect almost none of them to be homozygous, and if all of them were inherited from their sexual ancestors we would expect a third of them to be homozygous (see methods). We find that the mean proportion of homozygosity in asexual specific variants is 0.32 (Fig. 3D), which in the absence of recombination would suggest that the vast majority (97%) of asexual specific variants were inherited from sexual ancestors and then lost from the sexual population. This would correspond to only 914 de-novo synonymous mutations in the asexual population since the sexual-asexual split, which is unlikely given that the asexual population split from the sexuals hundreds of thousands to millions of years ago (Lázaro et al., 2011). Thus, we conclude that some form of recombination acts as a homogenising force.

However, we wanted to explicitly test whether variant inheritance from a sexual ancestor followed by subsequent loss in the sexual population can explain the homozygosity levels in uniquely asexual variants. To this end we assessed the proportion of homozygosity in uniquely asexual variants that are fixed within the asexual population. These represent the ancestral asexual variants of our population, and so are likely to be older than other variants that are only found in a subset of samples. We found that the mean proportion of homozygosity in ancestral asexual specific variants was 0.58, suggesting that there is excess homozygosity even if all these asexual specific ancestral variants were originally inherited from sexual ancestors rather than forming de-novo in the asexual lineage (Wilcoxon signed rank; Nancestral asexual specific = 2,275, V= 2,121,916, P < 0.001, Fig. 3E). Additionally, ancestral variants shared between the sexual and asexual population are more homozygous than asexual specific variants (Wilcoxon rank sum; Nasexual-ancestral = 9,206, NAsexual specific = 30,463, W = 211,577,670, P < 0.001, Fig. 3D), suggesting that more ancient variants are more likely to be homozygous. We also found that within asexual specific variants, those found in more samples were more likely to be homozygous (Binomial GLM, N = 30,463, P < 0.001, Fig3E). Taken together, this suggests that some homogenising process might be taking place in the asexual genome, and that its impact is larger on the more ancient variants compared to those with a more recent origin. These results indicate a genome wide mechanism in driving the loss of heterozygosity in asexual planarians that could potentially limit mutation accumulation in this asexual lineage.

### GC-biased gene conversion in asexual planarians

We hypothesised that since gene conversion has a GC bias (Chen et al., 2007), homozygous variants should display this signature if they were indeed generated by gene conversion. We find that homozygous variants are GC enriched compared to heterozygous variants, with 53% of homozygous variants being composed of GC nucleotides compared to 35% in heterozygous variants (Chi-sqr = 1,328.6, df = 1, NHOM = 17,487, NHET = 22,517, P < 0.001, Fig. 4A), suggesting a GC bias in the mechanism driving high asexual homozygosity.

**Figure 4.**
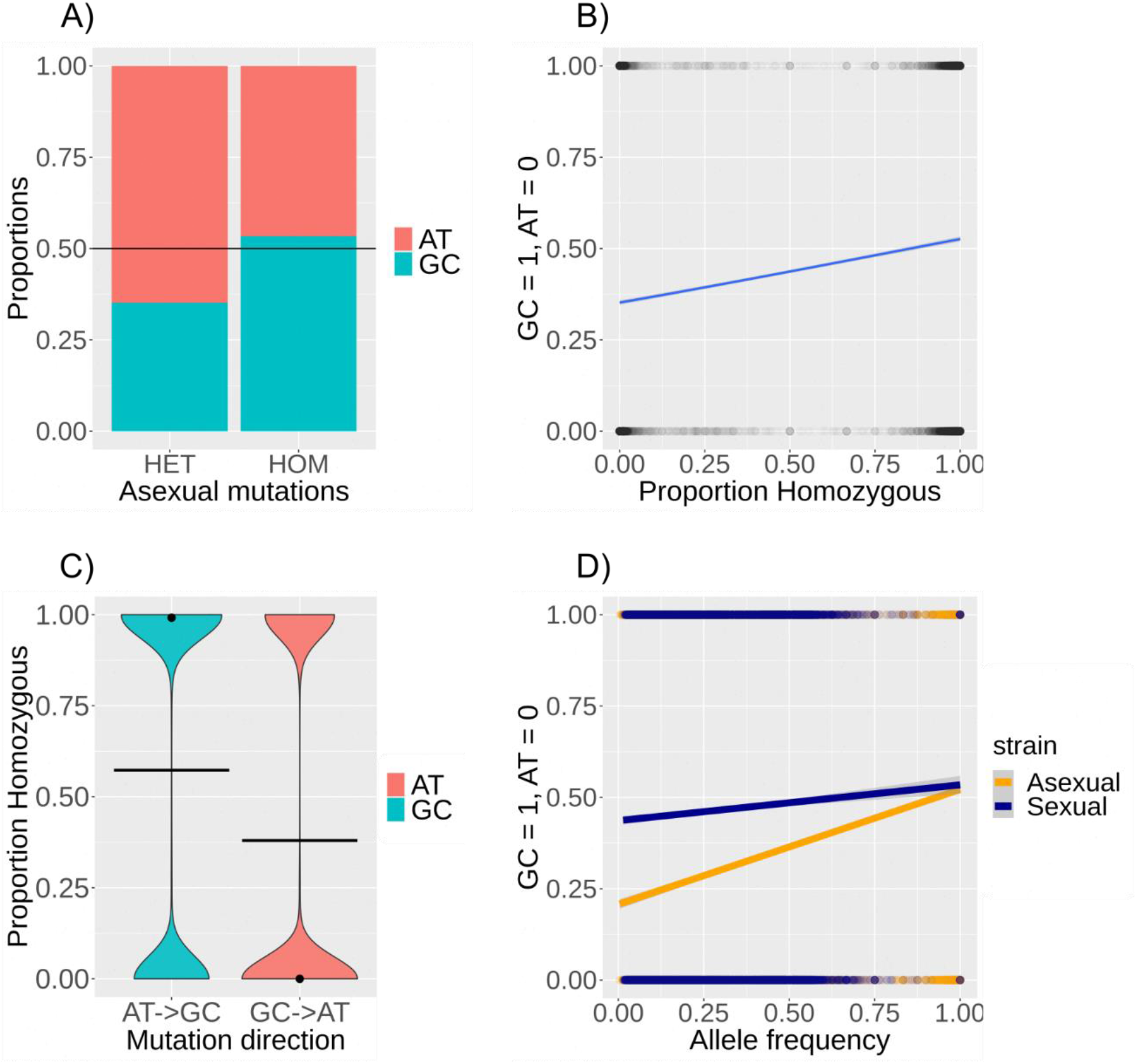
- GC bias in homozygous variants. A) Bar plot showing that the proportion of GC is higher in homozygous variants compared to heterozygous variants. Black line indicates the 50% line; B) Binomial GLM showing that variants with a higher proportion of homozygosity are more likely to be of GC identity in asexual planarians; C) Violin plot showing that AT to GC mutations are more homozygous than GC to AT mutations. Black dots indicate medians and black cross bars indicate means; D) Binomial GLM showing that variants with higher allele frequencies are more likely to be of GC identity in both sexuals and asexual, but in asexuals the effect is larger.

Additionally, we find that homozygosity is a significant predictor of GC-AT identity, with more homozygous variants being more likely to have a GC identity (Binomial GLM, N = 44,669, P < 0.001, Fig. 4B). Furthermore, we show that AT to GC mutations are 1.5 times more homozygous than GC to AT mutations, with mean homozygosity proportions of 0.57 and 0.38 respectively (Wilcoxon rank sum, NAT->GC = 17,984, NGC->AT = 21,483 W = 232,985,919, P < 0.001, Fig. 4C). This GC bias is not driven by AT to GC mutations across the transcriptome generally (Fig. S3). Finally, we find that variants with higher allele frequency are more likely to have a GC identity, and that this bias is stronger in the asexual strain compared to the sexual (binomial GLM, P < 0.001, Fig. 4D). Taken together, this suggests that GC biased gene conversion takes place in the asexual planarian, leading to a loss of heterozygosity which could limit the impact of Muller’s ratchet (Box. 2).

### Selection on gene conversion events removes deleterious mutations

Since gene conversion has the potential to generate homozygotes for both beneficial and deleterious variants, selection must be able to remove deleterious gene conversion events for gene conversion to be a viable solution to Muller’s ratchet. We find that mutations that are likely to be more deleterious due to their effect on predicted proteins (see methods), are less likely to be homozygous, suggesting that deleterious gene conversion events are selected against. We find that coding SNPs have a mean homozygosity proportion of 0.41 and are more homozygous than coding indels which have a mean homozygosity proportion of 0.25 (Wilcoxon rank sum; Nindels = 883, NSNPs = 64,220, W = 23,118,081, P < 0.001, Fig. 5A).

**Figure 5.**
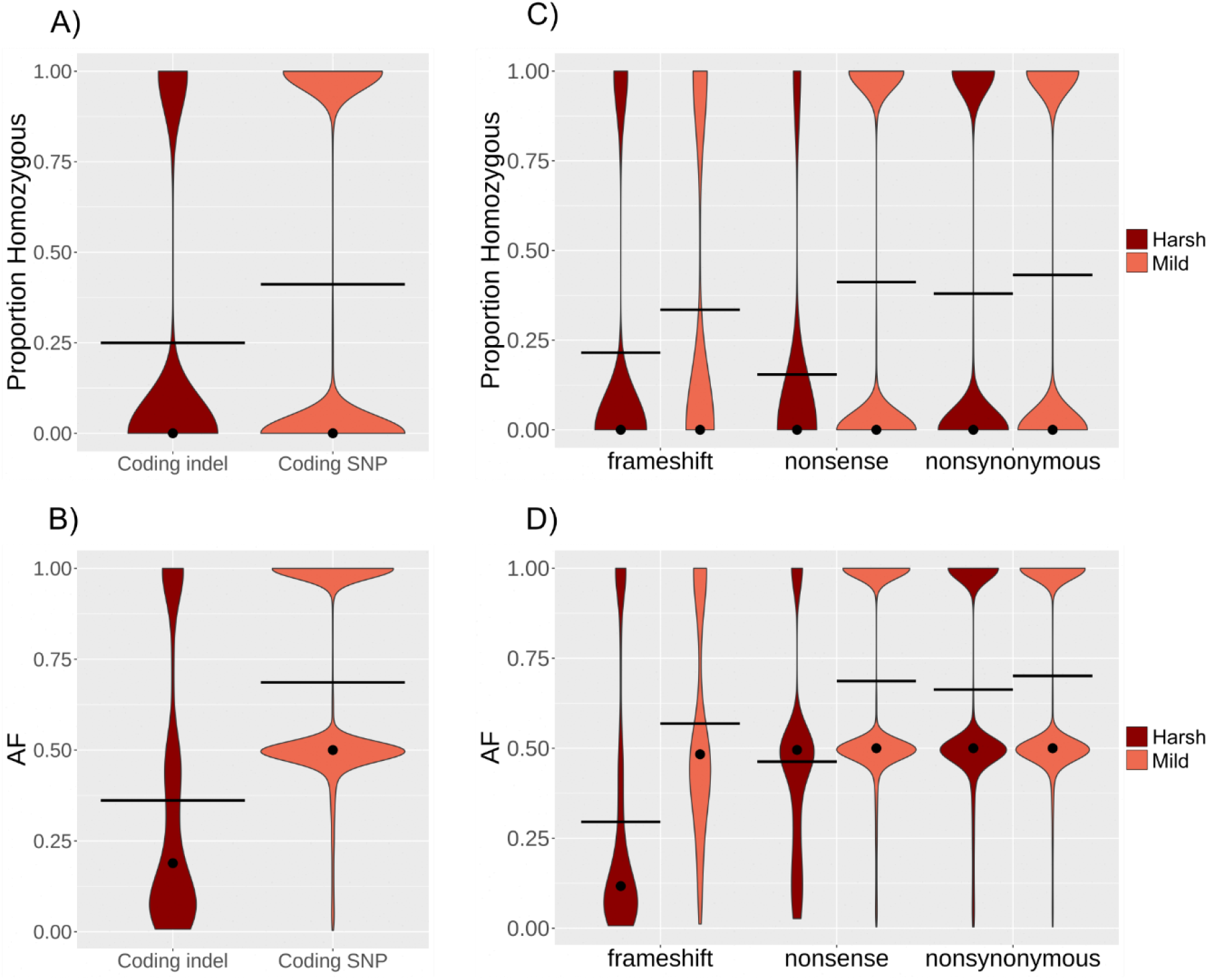
- Predicted deleterious mutations are less likely to be homozygous. Dots indicate medians, crossbars indicate means. A) Mean proportions of homozygosity are lower in coding indels than coding SNPs; B) Mean allele frequencies are lower in coding indels than coding SNPs; C) Frameshifting indels have lower homozygosity than non-frameshifting indels (mean proportion homozygous of 0.215 and 0.335 respectively). Nonsense variants have lower homozygosity than nonsynonymous variants (mean proportion homozygous of 0.15 and 0.38 respectively). Nonsynonymous variants have lower homozygosity than synonymous variants (mean proportion homozygous of 0.38 and 0.43 respectively); D) Frameshifting indels have lower allele frequencies than non-frameshifting indels (mean allele frequencies of 0.3 and 0.57 respectively). Nonsense variants have lower allele frequencies than nonsynonymous variants (mean allele frequencies of 0.46 and 0.66 respectively).

Additionally, coding SNPs have higher allele frequencies than coding indels, with mean allele frequencies of 0.69 and 0.36 respectively (Wilcoxon rank sum; Nindels = 883, NSNPs = 64,220, W = 11,652,733, P < 0.001, Fig. 5B). These patterns could be driven by other factors such as error rates in each category of mutation, so we also compared the homozygosity and allele frequencies of mutations within each category. We found that the homozygosity and allele frequencies were higher in synonymous vs nonsynonymous SNPs (Wilcoxon rank sum; NSynonymous = 39,669, NNonsynonymous 24,471=, WHom = 462,855,666, PHom < 0.001, WAF = 452,432,400, PAF < 0.001, Fig5C-D), non-framshifting indels vs framshiting indels (Wilcoxon rank sum; NNon-frameshifting = 154, NFrameshifting = 664, WHom = 44,439, PHom < 0.001, WAF = 23,168, PAF < 0.001, Fig5C-D), and Nonsynonymous SNPs vs nonsense SNPs (Wilcoxon rank sum; NNonsynonymous = 24,471, NNonsense = 65, WHom = 613,538, PHom < 0.001, WAF = 494097, PAF < 0.001 Fig. 5C-D). This suggests that mutations more likely to be deleterious are less likely to be homozygous and are rarer in the population. However, in contrast to this pattern, mutations in noncoding regions were altogether more heterozygous and had lower allele frequencies than those in coding regions (Wilcoxon rank sum; NNoncoding = 21,908, NCoding = 74,724, WHom = 891,850,288, PHom < 0.001, WAF = 860,143,632, PAF < 0.001, Fig. S4) That being said, the majority of noncoding mutations are found within repetitive regions (Fig. S4) which were shown in previous studies to have high mutation and erroneous mapping rates that may also explain their abnormally high levels of heterozygosity (Nishant et al., 2009; Patil and Vijay, 2021). We propose that when gene conversion events lead to a homozygous deleterious mutation in a cell, the homozygous deleterious clone is eliminated by intra-organismal and inter-organismal selection, leading to the lower observed homozygosity in variants that are predicted to be more deleterious. The separation of alleles via gene conversion followed by subsequent selection against unfit stem cell clones may help limit the impact of both Muller’s ratchet and the Hill-Robertson effect. Nonsynonymous variants have lower allele frequencies than synonymous variants (mean allele frequencies of 0.66 and 0.7 respectively).

## Discussion

Mechanisms explaining how some asexual species persist despite the potential costs of asexuality have remained elusive. Our results support a role for gene conversion in unlinking loci in the asexual genome of *S. mediteranea*. Combined with selection against deleterious homozygotes this allows these asexual planarians to limit the impact of Muller’s ratchet and Hill-Robertson effects, potentially explaining how this obligate asexual can persist over evolutionary time.

Our results suggest that the impact of Muller’s ratchet is relatively limited in asexual planarians. This is surprising since it has been shown both empirically and by simulation that Muller’s ratchet should drive the rapid accumulation of deleterious mutations even in recent asexual lineages (Loewe and Lamatsch, 2008; Lynch and Gabriel, 1990; Neiman et al., 2010; Tucker et al., 2013). For example, *Daphnia pulex*, an asexual lineage that is estimated to be only around 1,000 years old, is already showing signs of deleterious mutation accumulation (Tucker et al., 2013). Additionally, *Potamopyrgus antipodarum*, a freshwater snail in which the sexual form coexists with a recent obligate asexual form, shows higher rates of deleterious mutations in the asexual strain (Johnson and Howard, 2007; Neiman et al., 2010). This suggests that Muller’s ratchet can act rapidly to drive mutation accumulation in asexuals. Thus, one might expect asexual planarians to show clear signs of Muller’s ratchet, given that the asexual stem cells that make up their soma have been asexual for at least tens of millions of generations. However, they seem to only show an 27% increase of dN/dS in exons under negative selection, and this may be driven by the higher number of variants in the asexual dataset, increasing the statistical power for calling exons under negative selection and including those with higher dN/dS ratios. Furthermore, there is no evidence for elevated dN/dS ratios in highly conserved BUSCO genes, where nonsynonymous mutations are likely to be deleterious.

This lack of high mutation accumulation in asexual planarians could explain how they have persisted as asexuals and is similar to the patterns seen in some other long term asexual lineages (Brandt et al., 2017; Swanstrom et al., 2011). For example, asexual oribatid mites have been shown to have lower dN/dS ratios compared to their sexual counterparts, suggesting that, in this case, purifying selection might be stronger in the asexual species (Brandt et al., 2017). In bdelloid rotifers nuclear genes do not show signs of Mullers ratchet (Birky et al., 2005; Welch and Meselson, 2001). Others have shown an accumulation of deleterious mutations in bdelloid mitochondrial genes (Barraclough et al., 2007), but have later suggested that this may be due to other factors which when taken into account suggest no signs of Muller’s ratchet in the mitochondrial genes of bdelloid rotifers (Swanstrom et al., 2011). The lack of strong impacts of Muller’s ratchet on some long term asexuals suggests that they may possess mechanisms to limit the accumulation of deleterious mutations.

Some authors have previously suggested gene conversion as a mechanism to limit Muller’s ratchet in asexual lineages (Flot et al., 2013; Loewe and Lamatsch, 2008; Schön et al., 2009). The idea is that deleterious alleles can be overwritten by fit alleles, and that selection will favour these events, ultimately removing deleterious mutations from the population (Box. 2). However, beneficial alleles can equally be overwritten by deleterious alleles in the gene conversion process, and so selection must be able to act against these events to effectively counteract Muller’s ratchet. This might explain why some asexuals show high rates of gene conversion without limiting Mullers ratchet (Omilian et al., 2006; Tucker et al., 2013).

### Box. 2

Gene conversion as a method to remove deleterious mutations. Gene conversion can limit the progression of Muller’s ratchet by removing deleterious mutations from some of the progeny. A heterozygous diploid is depicted by homologous regions of red and blue chromosomes. The red chromosome contains a region with a deleterious mutation. The progeny containing a gene conversion event which removes the deleterious mutation in the red chromosome has the highest fitness (left) and can be selected for, while gene conversion events leading to a homozygous deleterious mutation (right) will be selected against. The most deleterious genome (a/a) is indicated by high transparency, followed by a/A, and A/A respectively.

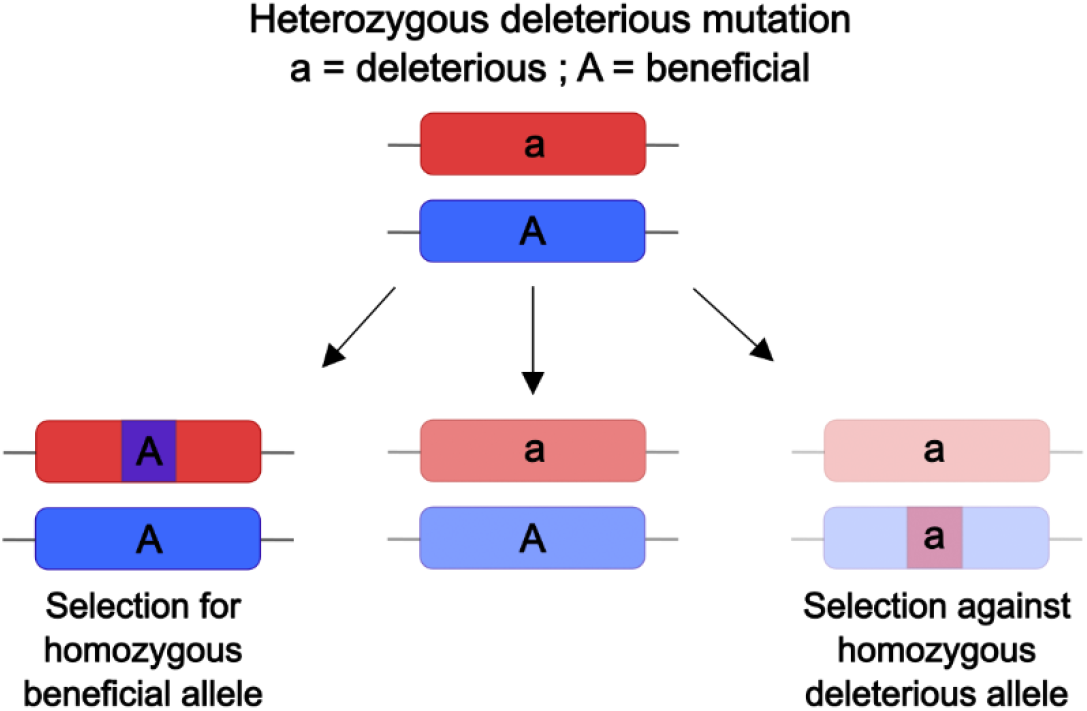

Furthermore, for gene conversion to stop the progression of Mullers ratchet, it may need to occur at a rate higher than the rate of point mutations. Otherwise, mutations may accumulate at a rate faster than gene conversion can remove them. This might be the case in some asexuals such as *Timema* stick insects, which show lower rates of gene conversion compared to their sexual relatives, and dN/dS values 3.6-13.4 times higher compared to their sexual counterparts (Bast et al., 2018; Henry et al., 2012). Despite these caveats, gene conversion remains a potential mechanism for limiting the cost of asexuality. This is in part because it is a byproduct of DNA damage repair mechanisms that are available to diploid (or higher ploidy) organisms (Chen et al., 2007) and so, it does not need to evolve *de novo* as a solution to asexuality. Supporting this are the shared signatures of gene conversion between bdelloid rotifers (Flot et al., 2013), darwinulid ostracods (Butlin et al., 1998), and now *S.mediterranea*.

While gene conversion may be important in both fissiparous and parthenogenic asexuality, we hypothesise that it might be significantly more effective in fissiparous asexuality. This is due to the variation within the soma being heritable in fissiparous organisms, increasing the amount of variation subject to selection (Howe et al., 2022). This is particularly relevant in *S.mediterranea* whose immortal stem cells are extremely abundant and represent a considerable proportion of the soma (van Wolfswinkel et al., 2014). These high numbers of stem cells are important since each stem cell will experience unique gene conversion events, and with a large enough population of stem cells it is likely that some of them will overwrite deleterious alleles with their beneficial counterparts. These stem cells may then outcompete other stem cells within the worm and spread, potentially reaching fixation in the worm.

Selection may then act between individual worms, selecting for those that have purged deleterious mutations, ultimately removing deleterious mutations from the population. Thus, fissiparous organisms may be more efficient than parthenogenetic asexuals at removing deleterious mutations from the population since gene conversion within their soma can also be selected upon to remove deleterious mutations.

However, it is important to point out that there is always the possibility that asexual planarians have rare cryptic sex that we are simply not aware of, a fundamental difficulty when researching the impacts of asexuality. For example, the brine shrimp, *Artemia parthenogenetica*, was once thought to be asexual but was later shown to undergo rare sex in a mass crossing experiment (Boyer et al., 2021). Additionally, even the presumed ancient asexual bdelloid rotifer *Macrotrachella quadricornifera* has had its asexuality contested with evidence supporting allele sharing between different isolates (Laine et al., 2022). However, similar patterns can potentially be explained without sex by ancestral duplications followed by independent allele losses (Wilson et al., 2024). Either way, it is entirely possible that in the case of *S. mediterrannea* rare cryptic sex might occur and might explain how they avoid the accumulation of deleterious mutations. However, it would not explain the strong signatures of gene conversion, which is not as clear in the genome of sexual planarians (Fig. 3C, Fig. 4D). Finally, even if rare sex occurs, gene conversion may still be useful as a mechanism to deal with the cost of highly reduced levels of sexual recombination.

In summary, obligate asexual reproduction in multicellular organisms is rare and thought to be an evolutionarily ephemeral phenomenon due to the fundamental limitations of asexuality. When we examine *S. mediterrannea*, which adopts this strategy successfully, we are able to assess how it limits the costs of obligate asexuality. Our results suggest that it likely uses asexual recombination in the form of gene conversion, to unlink loci and thus limit the impact of Muller’s ratchet and Hill-Robertson interference. While such mechanisms may be available to most asexual lineages, the opportunity for selection to act on a population of somatic stem cells may be key in allowing asexual planarians to persist.

### Materials and Methods Collecting available RNAseq data

RNAseq data was collected from previous studies with a total of 169 libraries from 7 studies for the asexual and 46 libraries from 3 different sexual populations (SI1) (Currie and Pearson, 2013; Davies et al., 2017; Duncan et al., 2015; Khan and Newmark, 2022; Scimone et al., 2014; Tu et al., 2015; van Wolfswinkel et al., 2014; Vogg et al., 2014; Zhu et al., 2015).

When choosing the studies, we only used those that used illumina sequencing and had clear annotations of the strains used. The Oxford sexual *S. mediterranea* population data was acquired by extracting RNA from whole, mature sexual worms using a standard Trizol extraction (Liu and Rink, 2018). Extracted RNA was then sequenced with Ilumina sequencing of 150bp paired end reads. Raw reads were deposited at the SRA repository (XXXX/XXXX).

### Variant calling and filtration

RNAseq data was then trimmed and quality filtered with trimmomatic and FASTQC using default settings (Bolger et al., 2014). STAR was then used to map the RNAseq reads to the dd_Smes_g4 genome (Dobin et al., 2013; Grohme et al., 2018). For variant calling we used the GATK 4.2.2.0 software following the best practice (Van der Auwera et al., 2013). We used the suggested parameters for variant filtration (SI9), except that we allowed for variants that had no reads supporting the reference (these are usually filtered out since some of the quality measures require a comparison of reads supporting the reference and alternative alleles). This provided us with a list of variants in each population (SI10).

### Defining Orthologues

We ran Orthofinder using a proteome from the Neiro et al annotation to isolate the orthologues of each predicted gene (Emms and Kelly, 2019; Neiro et al., 2022). To reduce redundancy, we first ran CD-hits to collapse all peptides with 99% identity to a single transcript ID.

### dN/dS analysis

We used the dNdScv R package for dN/dS analysis and annotation of mutation impact in coding regions (Martincorena et al., 2017). dNdScv uses an expansion on the Poisson framework to obtain dN/dS values and a full tri-nucleotide model to avoid systematic bias due to sequence context (Martincorena et al., 2017). To get the list of genes for dN/dS analysis we used the list included in the OrthoFinder output, and further filtered for the longest transcript per gene. We then ran dNdScv with no covariates and no limits on the number of variants per CDS. When comparing dN/dS values between the strains in genes under purifying selection, we used the local model from dNdScv, since the overdispersion value for the sexual dataset was lower than 1. To compare dN/dS values of BUSCO gene exons between the strains, we isolated single copy BUSCO genes in planarians using BUSCO (metazoan gene list). We then isolated BUSCO exons that had both synonymous and nonsynonymous mutations within them (to allow for meaningful dN/dS assessment using the dNdScv local model) and compared their dNdS values provided by dNdScv. We also compared dN/dS ratios in exons that were under negative selection in both strains. These were defined by having a qmis_loc < 0.05 and wmis_loc < 1 in the dN/dS output (SI3-SI4,

SI6). Finally, we used the global model to isolate genes under positive selection in asexual planarians (SI7-SI8), and top blast hits were taken from planmine (Rozanski et al., 2019).

### Analysis of zygosity and allele frequency

To assess allele frequency and homozygosity levels accurately, we only used information about variants from samples that had a minimum depth of 10 at the site (SI11). To achieve this, we split the VCF into individual VCFs (one per sample), and then filtered out any variants that were not covered by at least 10 reads. Allele frequency in each sample was calculated using VCFtools (0.1.16) freq function (Danecek et al., 2011). This resulted in each variant obtaining a 0, 0.5 or 1 value for each sample, with the numbers corresponding to homozygous reference, heterozygous, and homozygous alternative respectively. We could then use this information to calculate allele frequency in the population for each variant (total number of alternative alleles in the population / total number of alleles in the population). We also calculated the proportion of libraries that were homozygous for each variant (number of libraries homozygous for alternative variant / number of libraries containing the variant). Note that assuming a complete lack of recombination in asexuals, the measure of homozygosity from a group of individuals should be very similar to the measure of homozygosity within a single individual.

### Expected values for mean proportion of homozygosity

To test whether there is excess homozygosity in the asexual strain, we compared the observed and expected proportions of homozygosity in the asexual population. When testing for excess homozygosity, or GC bias in homozygous variants (Fig. 4), only synonymous mutations were used to limit the impact of selection. Assuming there is no recombination, the expected mean proportion of homozygosity for ancestral variants (those shared between sexual and asexual strains) is 0.5. This is because for a variant to be ancestral, it must have been inherited from a sexual worm as at least one of the alleles in the founding asexual population. Thus, whether an ancestral variant is homozygous depends only on whether the variants was also inherited as the second allele, meaning that half of the variants are expected to be homozygous. The expected proportion of homozygous asexual specific variants cannot be precisely known because it depends on how many of them were formed by mutation after the sexual-asexual split, and how many of them were inherited from a sexual ancestor and subsequently lost from the sexual population. However, we can place lower and upper limits for the expected homozygosity in the two extremes. The first case, which defines the lower limit, is a scenario in which all asexual specific variants arose by mutation after the sexual-asexual split. In the absence of recombination, we would expect this to lead to no homozygous variants (since the de-novo mutations should be heterozygous). In the second case, which sets the upper limit, all asexual specific variants were inherited from a sexual ancestor and so the proportion of homozygous variants depends on Mendelian principles. This means that there are 3 possible ways that the alternative alleles could have been inherited, alt/alt, alt/ref, and ref/alt, meaning that if all 3 options are equally likely, we would expect one third of all asexual specific variants to be homozygous (Fig. S4).

### Assessing evidence for effective selection on gene conversion events

To assess whether there is effective selection to remove deleterious gene conversion events, we wanted to test whether more harmful mutations are less likely to be homozygous. Since we cannot know the precise impact of each mutation on fitness, we broadly ranked variants based on the amount of change the mutation is likely to have on the protein sequence. In our analysis we assumed that coding indels are more deleterious on average than coding SNPs, frameshifting indels are more deleterious than non-frameshifting indels, nonsynonymous mutations are more deleterious than synonymous mutations, and nonsense mutations are more deleterious than nonsynonymous mutations.

It would also be reasonable to assume that that, on average, noncoding mutations are less deleterious than coding mutations. However, noncoding regions are often rich in repeats which can complicate the assessment of homozygosity and allele frequency. We wanted to test if this was indeed the case in our data, so we ran RepeatMasker 4.1.2-p1 on the dd_Smes_g4 genome which gave us a gff3 file containing all the repetitive regions in the genome. We then used bedtools (Quinlan and Hall, 2010) intersect to test the proportion of coding vs noncoding variants that are found within repetitive regions.

## Supplementary Data

SI1 – List of all libraries used in study (Sexual RNAseq from Oxford is unpublished). SI2 - Asexual dN/dS local model.

SI3 – Sexual dN/dS local model. SI4 – List of all exons used in study.

SI5 – List of all BUSCO exons with synonymous and nonsynonymous mutations in both sexual and asexual strains.

SI6 – List of all exons under negative selection in both sexual and asexual strains SI7 – Asexual dN/dS global model.

SI8 - List of exons under positive selection in asexual strain using the global dNdScv model.

SI9 – variant filtration criteria. S10 – List of variants

S11 – List of variants where homozygosity and allele frequency information only comes from libraries with at least a depth of 10 at the variant site.

## Acknowledgements

We would like to thank Tim Barraclough, Chris Wilson, and Tymoteusz Pieszko for their helpful comments on the manuscript.

## Data availability

Supplementary information and data can be found at: https://github.com/SimonOX-Gene/Smed_GeneConversion

Additional information is available upon request.

