## Supplementary material for "Gene conversion limits the cost of asexuality in somatically immortal worms": Supp_figures_010724

Supplementary figures

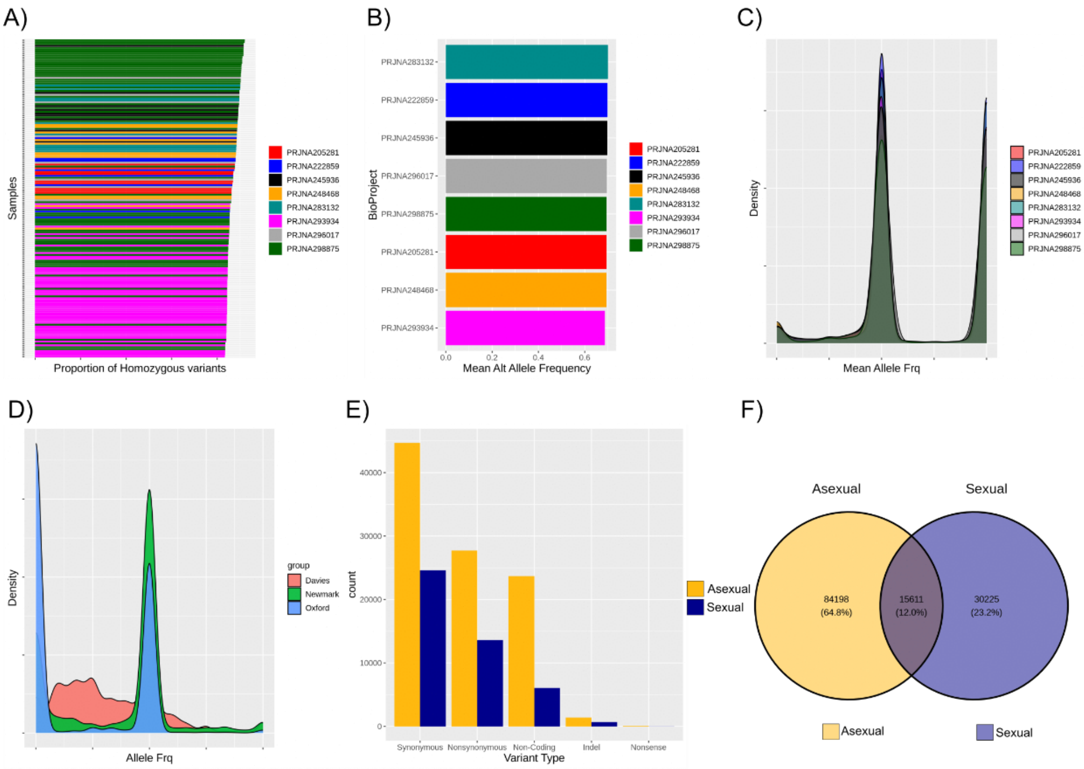


**Supplementary figure 1 –** Homozygosity and allele frequency across libraries. A) Proportion of homozygous variants in asexual libraries by BioProject; B) Mean allele frequency per BioProject; C) Distribution of allele frequencies in asexual BioProjects seem broadly similar; D) Distribution of allele frequencies in sexual populations. The sexual population from the Davies et al 2017 seems to have an abundance of rare alleles; E) Number of variants with informative homozygosity and allele frequency. These are variants where at least one of the samples covers with the locus with a depth of at least 10 (SI11). F) Venn diagram of asexual and asexual variants after filtering for a minimum depth per library (SI11)

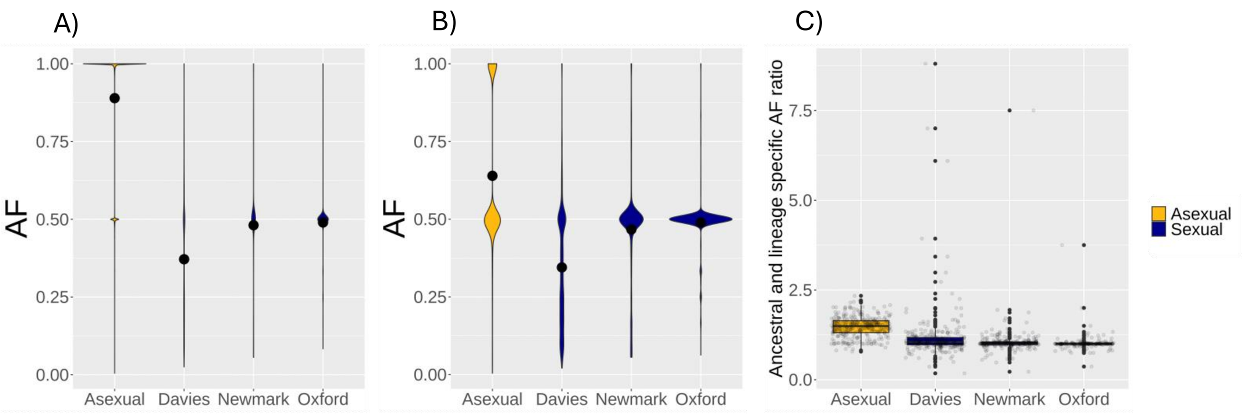


**Supplementary figure 2** – Allele frequency comparison with the sexual populations separated. A) Ancestral variants have higher allele frequencies in the asexual compared to each of the sexual populations populations (Wilcoxon rank sum; $N_{Asexual}$ = 14,206 , $N_{Daviesl}$ = 9,625, $N_{Newmark}$ = 7,898, $N_{Oxford}$ = 4,368, $W_{Asexua\text{-}Daviesl}$ = 126,240,405, $W_{Asexual\text{-}Newmark}$ = 97,403,170, $W_{Asexual\text{-}Oxford}$ = 53,920,171, $P < 0.001$). Note that they don't all have the same number of shared ancestral SNPs because they have different numbers of SNPs supported by sufficient depth (see methods); B) Lineage specific variants have higher allele frequencies in the asexual compared to each of the sexual populations (Wilcoxon rank sum; $N_{Asexual}$ = 30,463, $N_{Daviesl}$ = 11,438 , $N_{Newmark}$ = 8,978, $N_{Oxford}$ = 5,049, $W_{Asexua\text{-}Daviesl}$ = 277,930,015, $W_{Asexual\text{-}Newmark}$ = 165,802,180 , $W_{Asexual\text{-}Oxford}$ = 86,457,384, $P < 0.001$); C) Average allele frequencies per scaffold of ancestral vs lineage specific variants (split sexual groups) (Wilcoxon rank sum; $N_{Asexual}$ = 363, $N_{Daviesl}$ = 337 , $N_{Newmark}$ = 332, $N_{Oxford}$ = 246, $W_{Asexua\text{-}Daviesl}$ = 82,516, $W_{Asexual\text{-}Newmark}$ = 86,862 , $W_{Asexual\text{-}Oxford}$ =63,322, $P < 0.001$).

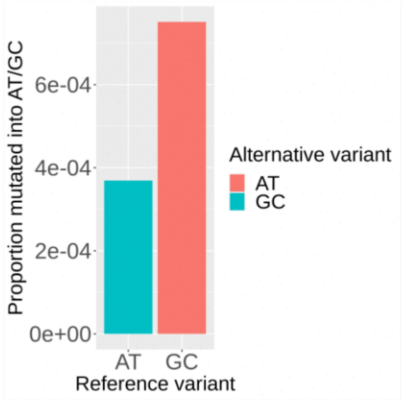


**Supplementary figure 3** – Proportion of AT variants that mutate into GC and vice versa using synonymous variants. Calculated by the number of synonymous AT to GC mutations divided by the total number of AT nucleotides in CDS regions (and vice versa for GC to AT mutations). This shows that GC nucleotides are more likely to mutate into AT nucleotides in the coding regions of the planarian genome, meaning that AT to GC mutations cannot explain the GC bias in homozygosity.

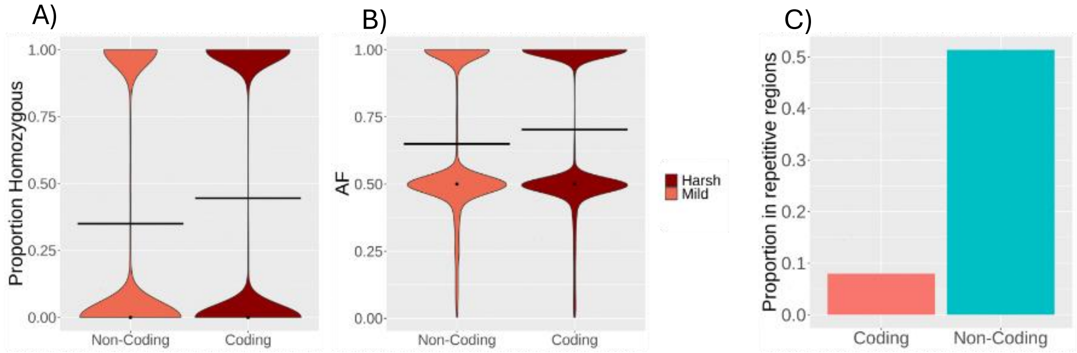


**Supplementary figure** 4 – Noncoding variants are more heterozygous and enriched in repetitive regions. A) Noncoding variants have a lower proportion of homozygosity than coding variants (Wilcoxon rank sum; $N_{coding}$ = 74,724, $N_{Noncoding}$ = 21,908, W = 891,850,288, P < 0.001); B) Noncoding variants have lower allele frequency than coding variants (Wilcoxon rank sum; $N_{coding}$ = 74,724, $N_{Noncoding}$ = 21,908, W = 860,143,632, P < 0.001; C) The majority of noncoding variants are found within repetitive regions, which might explain their high heterozygosity.

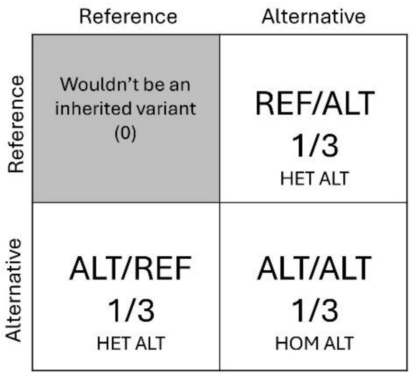


**Supplementary figure 5** – Punnet square explaining the expected proportion of homozygosity assuming all lineage specific asexual variants were inherited from sexual worms only to be subsequently lost in the sexual population. One third of the potential variants would be expected to be inherited as homozygous.
